# Volumetric imaging of an intact organism by a distributed molecular network

**DOI:** 10.1101/2023.08.11.553025

**Authors:** Nianchao Qian, Joshua A. Weinstein

## Abstract

Lymphatic, nervous, and tumoral tissues, among others, exhibit physiology that emerges from three-dimensional interactions between genetically unique cells. A technology capable of volumetrically imaging transcriptomes, genotypes, and morphologies in a single de novo measurement would therefore provide a critical view into the biological complexity of living systems. Here we achieve this by extending DNA microscopy, an imaging modality that encodes a spatio-genetic map of a specimen via a massive distributed network of DNA molecules inside it, to three dimensions and multiple length scales in developing zebrafish embryos.

---

Genomic mosaicism, the property whereby nucleotide-level differences present across the genomes of cells within tissues, are critical to organism biology and human health^1^. Data sets that have highlighted this intra-organismal and intra-tissue genomic diversity from the immune system^2^ to the nervous system^3^ have showcased the magnitude of missing detail in coarse gene-counts when they are not read out with DNA and RNA sequences from the same specimens.

These observations have therefore drawn a line between tissue genomic biology that is amenable to probe hybridization measurements^4^, where a gene’s status may be reduced to presence-versus-absence, and “de novo” sequencing-based assays which access a different level of information entirely. Concerted efforts to mend this blind spot include 2D biological pixelation^5^ to assign positional markers to DNA and RNA sequences, and sequencers built around individual samples^6^. In each of these cases, trade-offs between depth of focus, depth of capture, signal density, and resolution have placed hard bounds on the detail accessible. The genetic landscape of cell microenvironments – fundamentally phenomena that involve genetically unique three-dimensional neighborhoods – remains something we can only extrapolate, not image.

DNA microscopy is a distinct imaging modality that encodes an image of a single, “idiosyncratic” specimen into DNA using a stand-alone chemical reaction. It has previously been demonstrated in dense 2D multicellular specimens^7^ and has more recently been applied to the study of cell-surface protein polarity^8^. Theoretical variants have also been proposed^9;10^.

DNA microscopy begins by randomly tagging biomolecules inside a specimen with unique DNA-molecular identifiers, or UMIs. It then converts these DNA tags into an intercommunicating molecular network, where molecular copies of the original products are allowed to migrate, either by constrained or unconstrained diffusion, and link up.

The resulting linking frequencies encode spatial proximities of the original UMI tags, in the form of a UEI (unique linking-event identifier) matrix – whose rows and columns are individual UMI-tagged molecules. A statistical inverse problem is then solved on this matrix to infer the relative coordinates of the original UMIs. Any DNA or RNA sequence that these UMIs tagged may then be mapped to their corresponding locations, thereby assembling a complete spatio-genetic image of the original specimen.

Because DNA microscopy captures images from within a specimen and provides nucleotide-level readouts, it potentiates fully volumetric *de novo* (or zero-prior knowledge) spatio-genetic imaging. Two key barriers to broad application of DNA microscopy in three-dimensional tissues have been (1) high temperature thermal cycling for *in situ* PCR, that complicates uniform diffusion within the specimen, and (2) the separation of length scales that would need to be bridged computationally in order for a network of intercommunicating molecules/UMIs to inform the reconstruction of gross morphology.

Here, we overcome these barriers first experimentally, by introducing layered *in situ* chemistries that encode – at low and constant temperature – multiple length scales simultaneously into the output of DNA microscopy reactions. Second, we introduce an inference methodology that reconstructs encoded molecular positions over multiple length scales. We demonstrate its effectiveness on both earlier and newer data sets.

## Results

Encoding multiple length scales into a DNA microscopy data set requires engineering how UEIs either localize or de-localize from their UMIs of origin. We reasoned that a separation of UEI length scales could be achieved by initially dispersing UMI copies over short (<1μm) lengths via constrained diffusion and later dispersing UMI copies over long (>10μm) lengths via unconstrained diffusion. DNA microscopy^7^ had previously achieved the larger of the two length scales, using biocompatible PEG hydrogels formed around the sample to eliminate convection and limit the range of DNA molecule migration during the reaction to ∼50μm diameters^11^. We sought to execute this in parallel with ∼1μm diameter DNA dispersion^12;6^ achieved by rolling circle amplification, or RCA, in which the leading end of a DNA molecule, polymerizing along a circular template, diffuses while anchored to its point of origin.

### Volumetric DNA microscopy

The multiscale-encoding reaction is depicted in Figs 1 and S1. Briefly, RNA molecules in fixed cells were reverse-transcribed using random primers into cDNA, and 3’ DNA overhangs were added (Fig 1A). Pre-circularized DNA molecules, containing ∼25nt randomized UMI sequences, were then annealed to the ends of these protruding adapters (Fig 1B). Like in the first demonstration of DNA microscopy^7^, two distinct UMI types (“type I” and “type II”) for purposes of preventing homo-dimerization (the pairing of a UMI with itself) later in the experiment. A strand-displacing DNA polymerase elongated the annealed DNA polymer by RCA to create DNA nanoballs with tandem copies of the same UMI.

**Figure 1:**
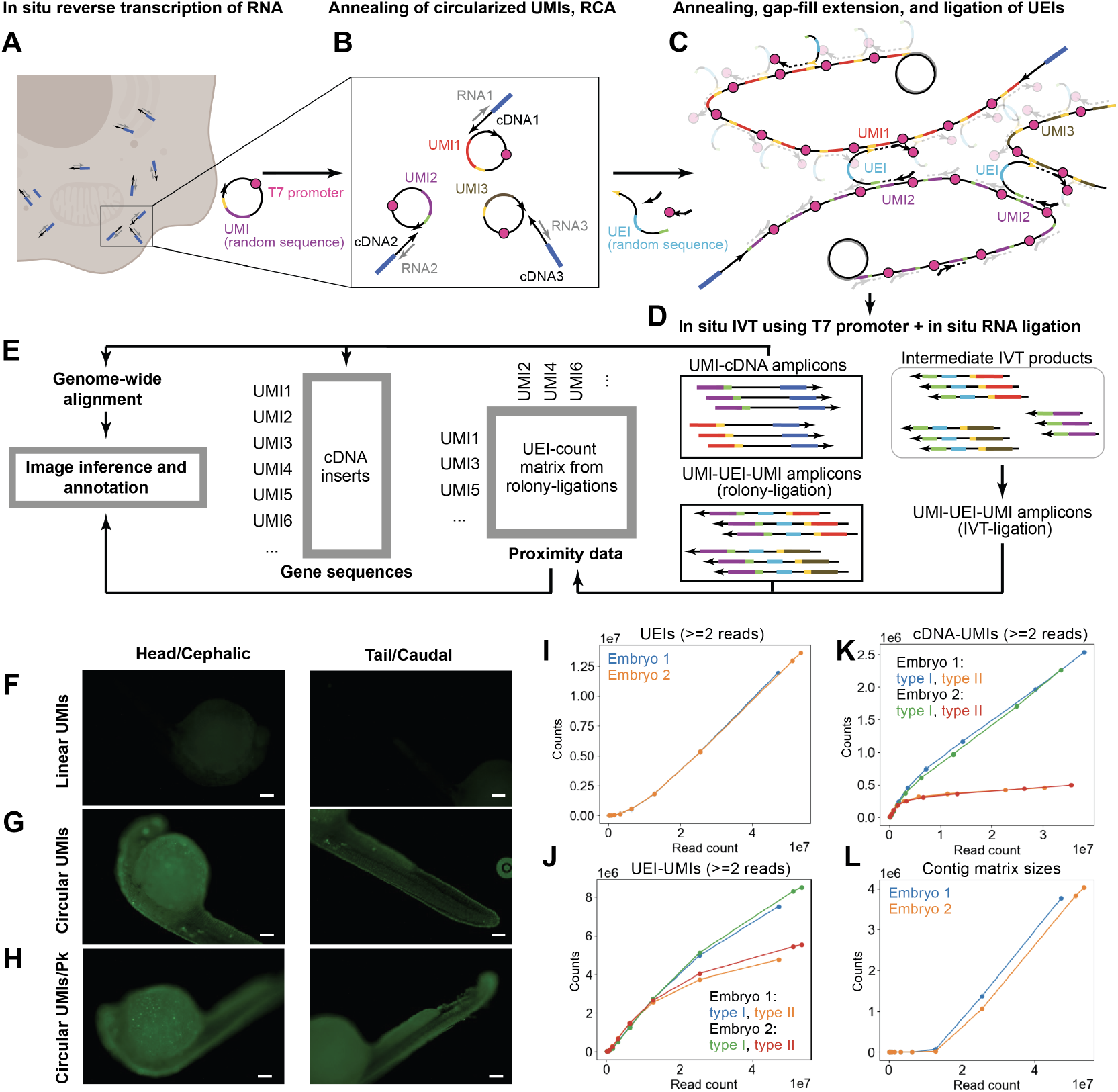
Volumetric DNA microscopy chemistry. begins by *in situ* synthesis of cDNA amplicons in fixed and permeabilized tissue (**A**). Pre-circularized UMIs are then added (**B**) to undergo RCA (**C**), which generates tandem copies of UMIs that undergo constrained diffusion about their points of origin. Oligos bridge adjacent UMIs to a new “UEI” amplicon. cDNA and UEI amplicons together undergo *in situ* amplification via in vitro transcription (**D**), with a complementary ligation reaction simultaneously fusing UMI-containing by-products into a different set of UEI-links. These data collectively encode proximity and cDNA sequence data (**E**). Using fluorescent nucleotides during RCA shows a lack of signal when linear UMIs are used (**F**) compared to circularized UMIs in zebrafish embryos permeabilized by methanol alone (**G**) or with proteinase K (**H**, scale bars 100*μ*m). Sequencing rarefaction of UEIs (**I**), UMIs from UEI amplicons (**J**), UMIs from cDNA amplicons (**K**), and contiguous UEI-matrix sizes (**L**) are shown.

The resulting DNA nanoballs physically pressed up against one another (Fig 1C), and a flanking oligonucleotide containing a randomized UEI sequence was then used to copy and label each UMI-UMI pairing uniquely. A combined in vitro transcription and ligation (“IVT-ligation”) reaction then amplified and further dimerized the resulting products (Fig 1D). All of these – abbreviated RCA-UEIs (UEIs generated from nanoball-adjacency), IVT-UEIs (UEIs generated from IVT-ligation), and cDNA (cDNA-UMI pairs) – were then further amplified by RT-PCR and sequenced (Fig 1E) for image inference and genome alignment (Table S1).

We first sought to determine whether RCA polonies in whole mount zebrafish embryos. Zebrafish embryos at 24 hpf were subjected to volumetric DNA microscopy chemistry (*SI: Experimental method*). We compared RCA reactions that incorporated fluorescent dUTPs by annealing either linear (Fig 1F) or pre-circularized (Fig 1G) UMIs. We found, as expected, DNA products generated in the latter but not the former. This signal was increased further by additional proteinase permeabilization of embryos (*SI: Experimental method*, Fig 1H). This demonstrated the permeability of the embryo under fixation conditions to circularized UMIs, enzymes, and other reagents.

Next, we performed “end-to-end” *in situ* reactions on 24 hpf embryos. The resulting number of distinct UEIs (Fig 1I), accompanying UMIs (Fig 1J), and separately amplified UMIs on cDNA-amplicons (Fig 1K) could be found increasing with read-counts (the latter preferentially amplifying with type-I UMIs). The sizes of contiguous UEI-matrices (Fig 1L), describing the number of UMIs that – through some set of UEI-links – were mutually connected, also increased with read-depth. Consensus cDNA amplicons were then mapped to the zebrafish genome (*SI: Sequence analysis*).

### Image inference

Inferring a DNA microscopy image of molecules (simulated positions separated by a UEI-association “fall-off” length scale: Figs 2A-B) from sequencing data is, at its greatest level of generality, a problem of calculating which putative molecular positions minimize a statistical distance (referred to as the “distance objective”) between UEI counts we expect given these positions and the UEI counts we observe (Fig 2C). This operation is mathematically equivalent to maximizing the probability of observations given these distances (*SI: GSE*, Figs S3, S4). Absent constraints, the solution to this problem is prone to both measurement noise and non-uniqueness.

**Figure 2:**
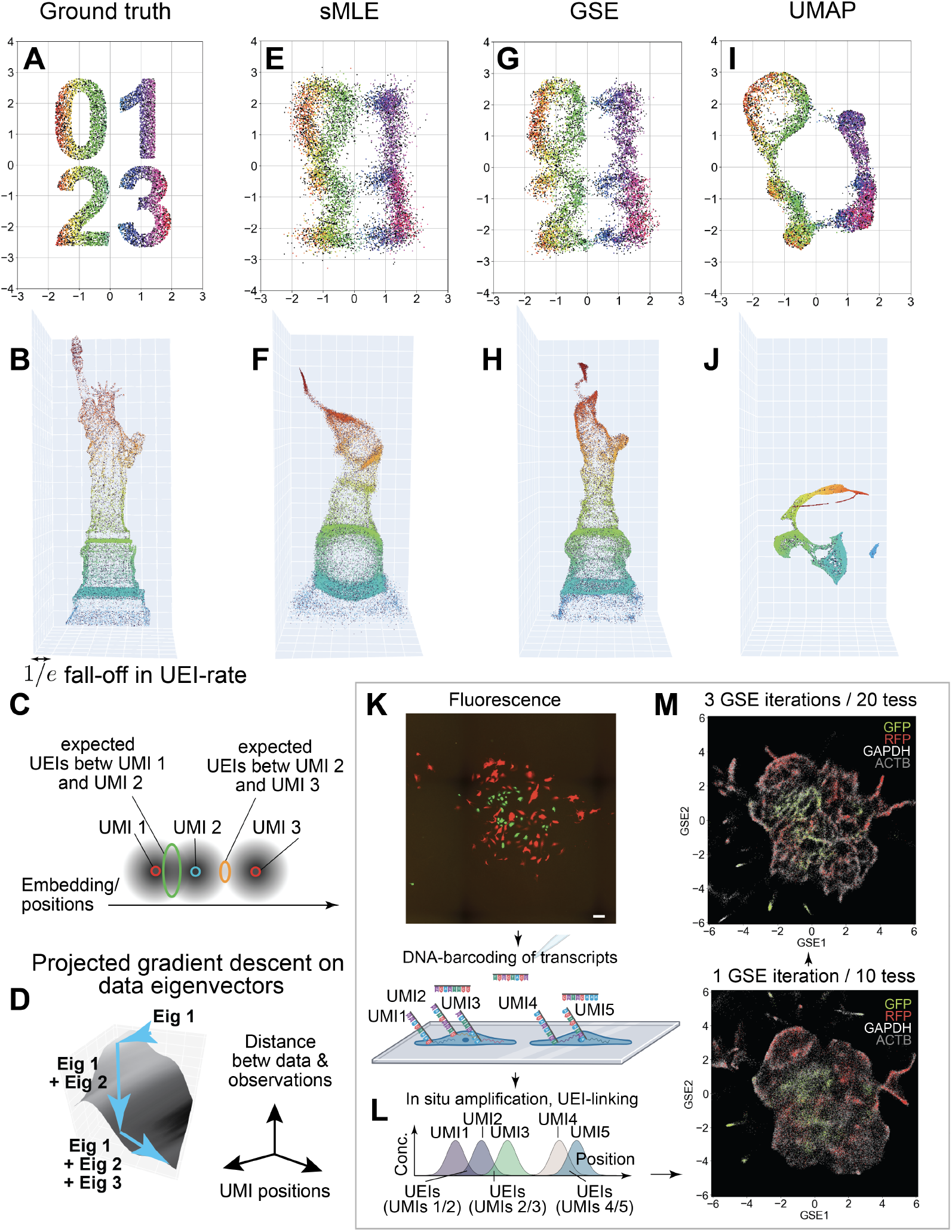
Geodesic Spectral Embeddings (GSE) applied to DNA microscopy simulation and experiment. Ground truth positions (**A-B**) are used to simulate UEI-count matrices that sample the distribution of pairwise proximities of points across the data set (1*×*10^4^ UMIs/1*×*10^5^ UEIs in 2D; 5*×*10^4^ UMIs/2.5*×*10^6^ UEIs in 3D). Image inference identifies UMI positions producing expected UEI counts that match those observed (**C**). Constraining these positions using data matrix eigenvectors (**D**) generates sMLE solutions (**E-F**). Modifying these eigenvectors with the GSE algorithm improves solutions (**G-H**). By comparison, UMAP alone produces distortions while obscuring geometry (**I-J**). Applying GSE to previous (6.5*×*10^4^ UMIs/7.9*×*10^5^ UEIs) DNA microscopy samples (**K**, scale bar 100*μ*m) whereby UMI-barcoded transcripts undergo unconstrained diffusion (**L**) allows sharpening of resulting inferred images (**M**).

Spectral maximum likelihood estimation, or sMLE, confines the position-solution to a linear combination of the top principal components, or eigenvectors, of the UEI matrix, analogous to a low-pass noise filter in optical imaging^7^. The optimal linear combination is determined by incrementally adding d eigenvectors (for a d-dimensional inference) to this set of eigenvectors in a “projected” gradient descent on the distance-objective until the solution converges (Fig 2D).

In two- (Fig 2E) and three-dimensions (Fig 2F), these solutions produce good but blurred approximations to underlying molecular coordinates. The reason for the shortfall of sMLE is that it uses the top eigenvectors of the “raw” UEI data matrix, which are solutions to a least-squares problem that weights all UEI counts equally.

In order to generate eigenvectors that account for differences in UEI-length scales and UMI-density across the data set, we developed a dimensionality reduction approach, called Geodesic Spectral Embeddings, or GSE. GSE directly approximates long-range curvature in the underlying manifold swept out by the data matrix in high-dimensional space. This is achieved by forming a kernel proximity matrix that describes not direct distances, but the “shortest traversable” distances along local linear tangents and their meeting points (*SI: GSE*). GSE eigenvectors are then used in precisely the same that raw-data eigenvectors are used in sMLE: as a basis for projected gradient descent of the full DNA microscopy solution.

Deriving GSE eigenvectors requires two parameter choices (*SI: GSE*). One is the degree to which the data is tessellated in order to analyze local “neighborhoods” of the UEI matrix. The second is the number of eigenvectors generated from the raw count matrix to analyze the curvature of the data manifold. Unless otherwise indicated, in the data shown we use 10 tessellations and 50 eigenvectors in each data set.

Applying GSE to simulated data (Figs 2G-H) significantly outperformed sMLE and UMAP^13^ (Figs 2I-J) in 2D and 3D. This demonstrated the algorithm’s generality in addressing non-uniform point distributions in higher dimensions.

We next investigated whether GSE could improve upon single length-scale DNA microscopy reconstructions^7^. In this earlier experiment, an ensemble of cells in culture is plated (Fig 2K, photograph from Weinstein et al 2019), and specified gene amplicons are tagged with UMIs (Fig 2L) that then undergo *in situ* amplification-reaction, unconstrained diffusion, and UEI-linking (analogous to the experimental design in Fig 1. The resulting UEIs are sequenced and the image is inferred. The fluorescent protein gene sequences read out from DNA microscopy are then compared back to the actual fluorescence measured in light microscopy of the same specimens.

Applying GSE to this data over a single iteration yielded resolution comparable to that found using sMLE previously^7^. Iterating GSE three times, each time treating newly generated GSE eigenvectors as updated “raw” count matrix eigenvectors (*SI: GSE*), as well as increasing the number of data-tessellations from 10 to 20, improved cell boundaries substantially (Fig 2M).

### Whole organism DNA microscopy inference

Having demonstrated GSE’s effectiveness on accurately reconstructing 2D DNA microscopy data sets and 3D simulations, we sought to evaluate its ability to reconstruct whole organism data sets, such as those described in Fig 1. An initial sub-sampling of 12.8 million reads from embryo 1 generated a 7*×*10^4^-UMI matrix (Fig 3A) with discernible structure. However, as the depth of sequencing increased and UMI count did so as well, the reconstruction blurred (Fig 3B), an artifact of the GSE algorithm over-fitting the “jaggedness” of the densely-populated manifold swept out by the more deeply sampled data (Fig 3C).

**Figure 3:**
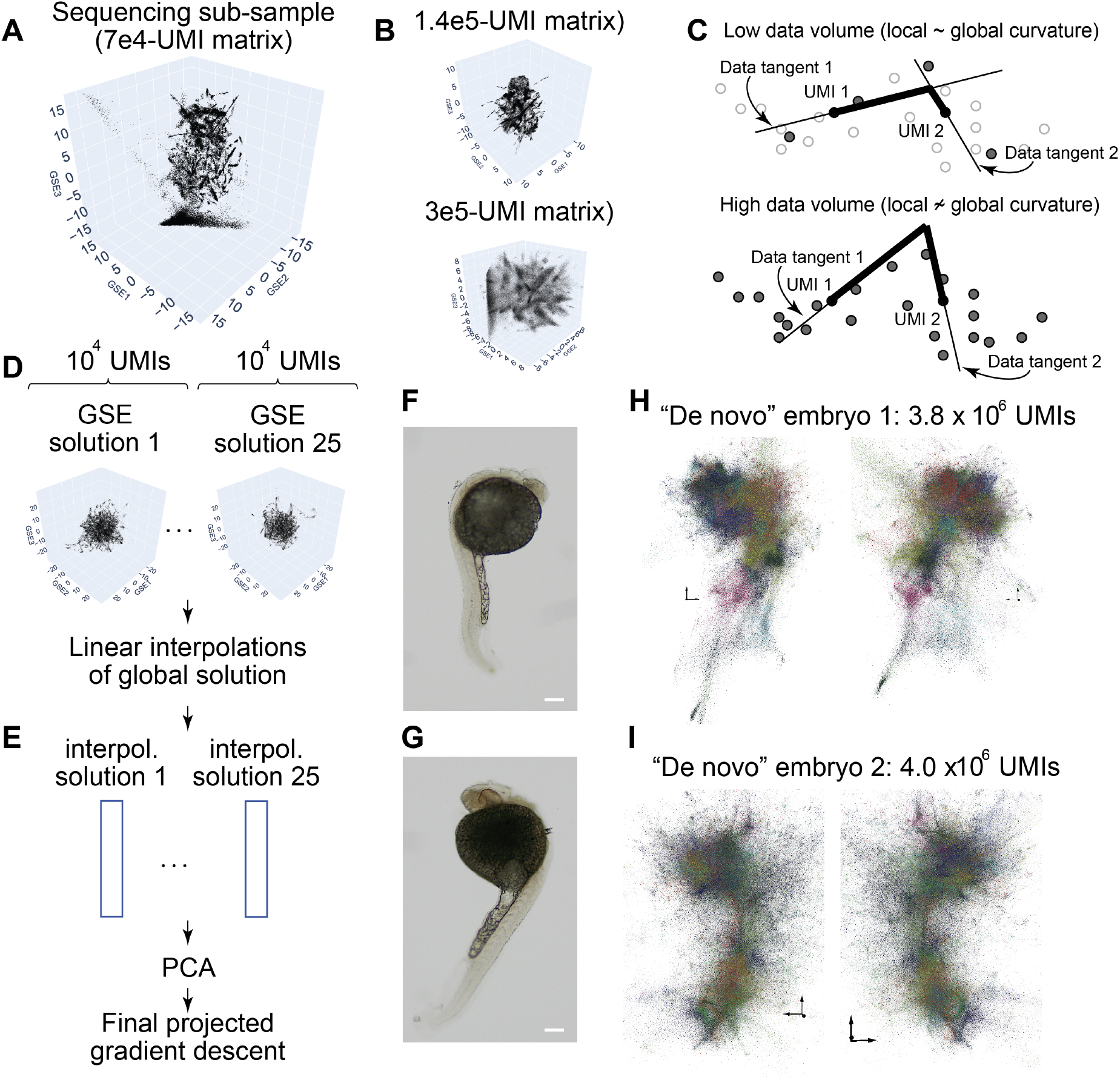
Large-scale 3D inference for whole-organism DNA microscopy. Low-depth sequencing and GSE-inference produces granular detail (**A**) that blurs at higher sequencing depth (**B**), which can be qualitatively explained by GSE’s sensitivity to data “jaggedness” on deeper sampling (**C**). To solve this, sub-sampled data-sets (∼10^4^ UMIs) generate granular “sub-solutions” (**D**) from which a putative reconstruction of all 10^6^ UMIs for each can be found by linear interpolation (**E**) and collated by PCA. UEI data from the original embryos 1 and 2 (**F** and **G**, respectively; scale bars 200*μ*m) are subjected to this image inference to give final GSE embeddings for the full 10^6^-UMI data sets (**H** and **I**, respectively, both showing two perspective angles of the same embryo). Distinct colors are arbitrarily assigned to distinct spectral clusters/segments. Axes in GSE plots indicate scales at different locations, with arrows having length of 3 GSE-units (1*/e*-association fall-offs).

To solve this, for each data set, we sub-sampled data to several small 10^4^ UMI data sets (Fig 3D). For each of these “sub-solutions” (here numbering 25), a linear interpolation was then performed to estimate the locations of all UMIs not included in that specific sub-solution (Fig 3E). Performing PCA on the UMI covariances then gave a new set of eigenvectors/components to construct the global solution by an incremental projected gradient descent, on the *original* probability function that modeled the DNA microscopy reaction, as before.

Taking the original embryos (Figs 3F-G), whole organism inferences were generated (Figs 3H-I). Spectral clustering (*SI: Clustering*) similar to that previously used for segmenting cells in DNA microscopy^7^ was applied to identify segments within the data, where a segment was defined as including UMIs that communicated discernibly more frequently with one another with others that were otherwise nearby. The resulting images, with colors introduced to visualize distinct clusters, recapitulated gross embryo morphology, highlighting a “lobed” structure to the head and a well-defined tail/caudal region. We next sought to examine the differential representations of gene transcripts across the inferred 3D images.

### Genomic sequence distribution analysis

We began by establishing a means to identify embryonic regions, by first aggregating UEIs between distinct segments colorized in Figs 3H-I, then taking the first eigenvector of the resulting UEI graph and bisecting points at its median. The resulting division of genome-mapped UMIs is depicted by distinct colors for embryos 1 and 2 in Fig 4A and B, respectively.

**Figure 4:**
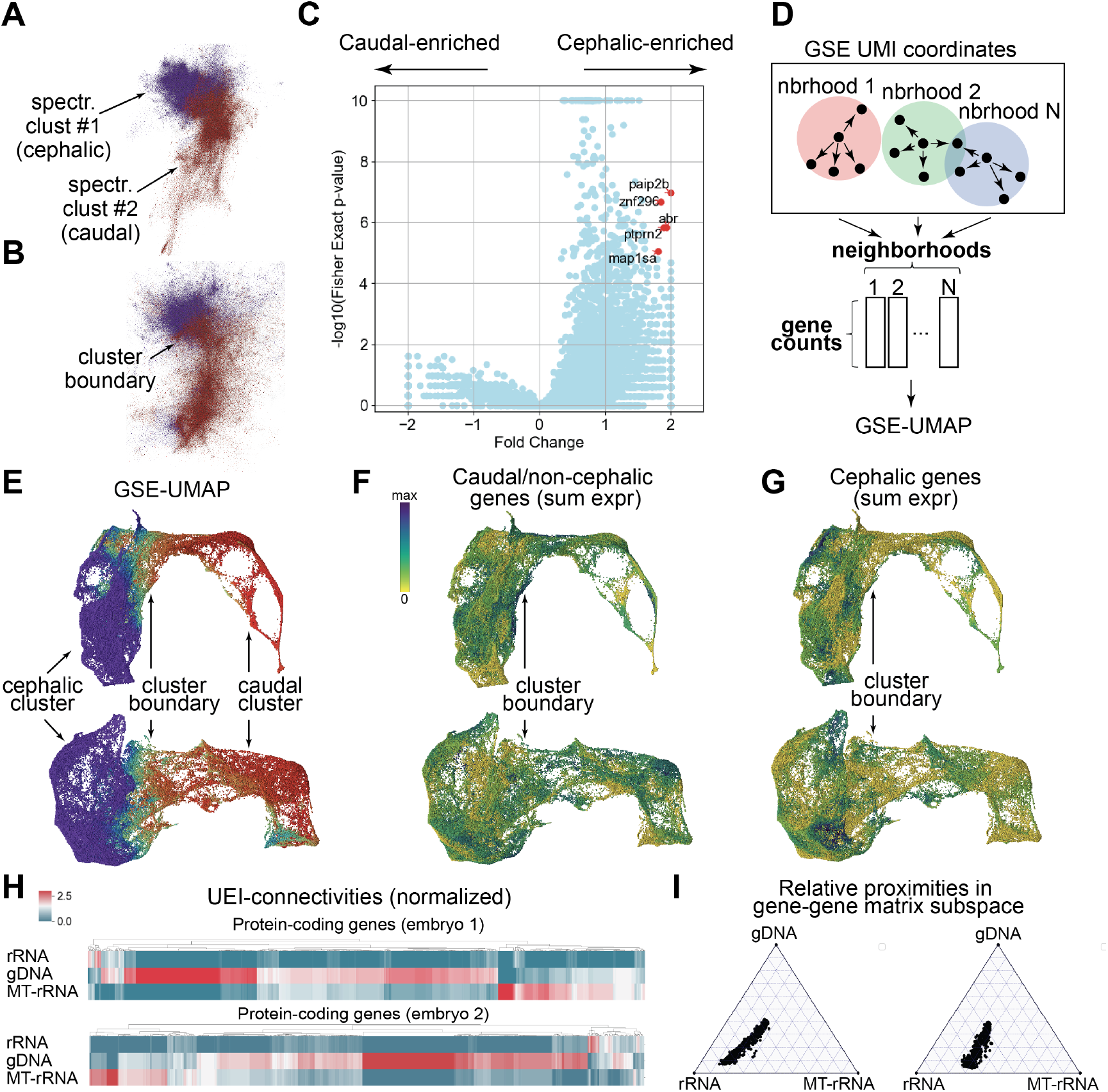
Spatio-genetic maps of embryos at multiple length scales. Spectral clustering allows embryos to be divided between cephalic/head and caudal/tail domains (**A-B**). Performing two-sided Fisher’s Exact Test on each mapped gene using this separation (fold-differences displayed relative to the averaged value) allows for the identification of gene enrichment (**C**). Performing UMAP-embedding on gene-counts in 10^5^ GSE-neighborhoods (**D**) provides a joint spatial-genetic embedding (**E**; color scale the same as in **A-B**) of embryos 1 (top) and 2 (bottom). Plotting a heat map on top of this embedding shows differences between summed expression of genes commonly expressed outside of the head region (**F**) and those predominantly expressed in the head region (**G**). The relative proximity of protein-coding genes to rRNA, MT-rRNA, and gDNA can be visualized either by direct UEI-connectivity, with genes shown as columns (**H**, only genes with *≥*10 UEIs summed across all three categories are shown; colors scaled as number of UEIs normalized to column mean), or by relative proximities in the subspace formed by the top eigenvectors of the gene-gene interaction matrix, with genes shown as points (**I**).

Next, we aggregated all genome-mapped UMIs together, and constructed contingency tables for each gene, that described the number of UMIs detected for that gene on each side of the median-location versus all other mapped UMIs. These contingency tables allowed us to calculate two-sided Fisher’s Exact Tests, each provided an effective upperbound on the p-values for the null hypothesis whereby all genes had the same distribution along the cephalic-caudal axis.

Although a large number of statistical differences were detected across the genome, we narrowed our focus to the five most head-enriched genes with p < 10^*−*5^ (Fig 4C). Of these five at 24hpf, two (znf296^14^ and ptprn2^15^) had been found exclusively in the brain by previous *in situ* hybridization (ISH) studies, one (map1sa^16^) had been found predominantly expressed in the head region and had a predicted role in neuronal physiology, and two (paip2b^16^ and abr^16^) were predicted more generally to be active in protein translation and cellular connections. Taken collectively, these were consistent with matching gene expression characteristics to the anatomical structures in the head and brain.

We next sought to highlight the spatio-genetic image’s recapitulation of known expression patterns still further by examining a distinct set of 20 genes commonly expressed in the head-region at 24hpf and 14 genes commonly expressed in the tail-region at 24hpf^16^ (Table S2). To do this, we implemented a joint spatial-genetic embedding scheme similar to a joint-embedding scheme previously used for immunoglobulins^2^.

Specifically, for each embryo, 1*×*10^5^ UMI “neighborhoods” were assigned by randomly choosing genome-mapped UMIs throughout the data set, and finding the 5000 nearest-neighbors of each (within the GSE embedding) that also mapped to the genome. These neighborhoods, sometimes overlapping (Fig 4D) formed a new high-dimensional data set (this time in gene expression space). The resulting vectors then underwent dimensionality reduction with UMAP (generated here using 100 PCs and 100 nearestneighbors).

The cephalic and caudal clusters of both embryos (Figs 4E, S2A-D) partitioned in a manner consistent with GSE alone (Figs 4A-B: same colorscale, with neighborhoods including UMIs from both cephalic and caudal clusters assigned an average color). Summing the expression of all 14 caudal (Fig 4F, Table S2) and all 20 cephalic (Fig 4G) genes drawn from previously measured ISH patterns yielded distinctive patterns consistent with the inferred locations along the caudal-cephalic axis.

Having established the recapitulation of gross morphology with DNA microscopy, we next sought to examine the application of such whole organism images to sub-cellular localization of molecular species. To do this, we summed UEI-counts between distinct genes/biotypes, rather than distinct UMIs.

The resulting graphs for embryos 1 and 2, depicted in relation to rRNA, MT-rRNA, and gDNA in Fig 4H, preferentially formed UEIs on a per-gene basis with gDNA, which was reduced when normalized to the UEI-sums of each of these three molecular species (Fig S2E-F). In the context of the full gene-gene connectivity matrix (using its top 100 eigenvectors, *SI: Clustering*), protein-coding genes showed closest proximity to rRNA (Fig 4I), consistent with their expected sub-cellular distribution.

## Discussion

We have demonstrated here the capability for a massive (>10^6^) distributed molecular network to volumetrically image a biological specimen from the “inside-out”. The implications of this work are threefold.

First, we have shown that DNA is capable of encoding massive images and that these images are capable of being decoded without the use of any specialized instrumentation beyond a DNA sequencer. This lays a critical foundation for the economy of scale this technology provides, and a broader democratization of 3D spatio-genetic imaging. This provides a clear path toward use in clinical settings, in which the impact of somatic mutation and genomic “idiosyncracy” in tumors^1^, lymphocytes^2^, the brain^3^ and the gut microbiome, play a critical role. Still further, the fact that all readouts are “zero-knowledge” opens up unexplored spatio-genetic complexity in non-model organisms.

Second, the inferred images here exhibit an inherent tension between being both connectivity maps and representations of Cartesian coordinates. We have shown that in simulation and practice, the implementation of a distinct methodology for dimensionality reduction – Geodesic Spectral Embeddings, or GSE – provides a scalable and reliable solution to reconciling these two disparate properties that is complementary to other common dimensionality-reduction methods. GSE may have broader application to other large data sets requiring similar reconciliation between the connectivities of nodes and their low-dimensional representations.

Third and finally, continued improvements to volumetric DNA microscopy chemistry and inference will provide a platform – distinct from and potentially complementary to conventional light and electron microscopy – for the analysis of biological circuits. The effective resolution of DNA microscopy follows a dependence on UEI-counts similar to stochastic super-resolution light microscopy’s dependence on photon-counts^7^, with the diffusion length scale (whether unconstrained, or constrained in the case of RCA) divided by the square root of the number of UEIs belonging to the resolved UMI. A 1μm size of RCA polonies with the 3 to 4 UEIs per UMI highlighted in Figure 1 therefore avails us of roughly the same length in resolution. Sequencing deeper, and increasing reaction yield would, however, push us well below the sub-micron regime.

In this work, we have demonstrated the ability for volumetric DNA microscopy to capture both RNA and DNA, and looking forward we anticipate acquiring proteomic details via oligo-antibody conjugates. As efforts accelerate to perform system-wide maps of neuronal circuitry in particular, the need to supplement these insights with those from molecular genetics, from gene expression, to genomic mosaicism, to spatio-proteomic measurement will take on increasing importance. We view volumetric DNA microscopy as poised to form a critical foundation for this broader undertaking.

## Acknowledgments

We thank John Kilkus and Lillian Roderick-Buescher for help with logistics, Dr. Andrew Piacitelli for early work not included in this study that has informed our ongoing research, and Dr. Victoria Prince for help procuring zebrafish specimens. We thank Dr. Raghu Mirmira and Dr. Ryan Anderson for helpful discussions, and Dr. Feng Zhang and Dr. Aviv Regev for guidance. This work was funded in part by the Damon Runyon Foundation, the Moore Foundation, NIH-1R35GM143017-01, NIH-5R01HG009276-04 (sub-K), and NSF-2121044. The authors have filed a patent application related to this work.

## Supplementary Information

### Data availability

Code and documentation can be found at https://github.com/wlab-bio/vdnamic. Raw sequencing data is available at SRA PRJNA1004618.

### Experimental Method

All reagents used are enumerated in Table S3 and all oligonucleotides are enumerated in Table S4.

#### Zebrafish embryos preparation

AB-wildtype zebrafish were kept and crossed in accordance with the approved protocols and ethical guidelines of the University of Chicago Institutional Animal Care and Use Committee. Embryos were collected at 24 hours post fertilization (hpf), dechorionated with 1mg/ml pronase 5min at 28C. Dechorionated embryos were fixed in 4% paraformaldehyde in 1xPBS at 4C overnight. After dehydration in 100% methanol for 15-30min at room temperature, the embryos were stored at -80C for at least 2hrs before use. Embryos were successively rehydrated with 75%, 50% and 25% methanol in 1xPBS for 5min each and washed 4x with 1xPBST (1xPBS + 0.1%Tween-20), 5min per wash at room temperature. The embryos were then permeabilized with 0.5-1 x 10^*−*4^ U/ul Thermolabile proteinase K for 12min at room temperature. The Proteinase K was then inactivated at 55C for 15min. Samples were then washed 4x in PBST, 5min per wash.

#### RT (Reverse transcription)

After permeabilization, embryos were incubated at 4C for 1hr under slow rotation (10rpm) followed by 10min of 65C incubation in a pre-RT buffer comprising 20% formamide, 0.5U/ul Superase-In, 4.4mM DTT, 0.5 ug/ul rBSA, in 1xPBS and then cooled down to 4C immediately. After one water rinse, reverse transcription mix (1x FS buffer, 4.4mM DTT, 400uM dNTP, 32uM aminoallyl-dUTP, 0.5ug/ul rBSA, 1U/ul Superase-In, 10U/ul Superscript III, and 1uM 21.068C-8N RT-primer) was added and underwent 4C incubation for 1hr, 60C 3min, and 37C overnight under slow horizontal orbital rotation, followed by 1hr of 50C incubation. After RT, embryos were washed 3x in PBST and 1x with water, incubated in ExoI mix (1x ExoI buffer, 1.43U/ul ExoI) at 4C 1hr under slow rotation, followed by 37C 1hr to remove the RT primer and displaced cDNA. Embryos were again washed 3x in PBST and 1x with water.

#### Tagmentation

Transposomes were assembled according to manufacturer protocol. Briefly, oligos 22tn5.003A and 22tn5.MOS-3p were resuspended to 100uM individually in annealing buffer (40mM Tris-HCl pH8, 50mM NaCl). These were annealed at equal molar ratio at 95C 2min, followed by 1 degree-C decrements per minute for 70min. Transposome assembly mix (25uM of the annealed oligo-duplex, 1ug/ul tagmentase) was then mixed and incubated at 23C for 30min. Glycerol was then added to 50% and stored until use at -20C. Following reverse-transcription of samples transposome-glycerol stock was diluted 3:10 in tagmentase dilution buffer. This diluted solution was then added at a further 1:50 dilution to transposome reaction mix containing 5mM MgCl_2_, 10mM Tris-HCl (pH7.5), 10% N,N-Dimethylformamide, 9% PEG8000, and 850uM ATP. This mix was added to samples, and incubated at 4C 1hr under slow rotation followed by 55C 1hr. Samples were then washed 2x in PBST and 1x in PBS.

#### Cross-linking and transposome denaturation

BS(PEG)5 mix was prepared in 1xPBS to 5mM concentration and added to samples for incubation at room temperature for 1hr under slow rotation. Samples were then rinsed with 1M Tris pH 8, and quenched in this buffer 30min at room temperature. 4xSSC was then added, samples were incubated at 4C for 10min under slow rotation, and then at 70C for 15min. 10% formamide/2xSSC was then added, the samples were incubated at 4C under slow rotation for 10min, and then incubated at 50C for 10min. Samples were then washed 3x in PBST, 5min each.

#### 3’ adapter ligation and circDNA annealing

After rinsing with water, 3’ adapter ligation mix (500nM 22tn5.005, 500nM 22tn5.006, 1.25U/ul SplintR ligase, 1x SplintR Reaction buffer containing ATP) was added to samples, which were incubated at 4C 1hr followed by 23C overnight. After ligation, samples were washed 3x in PBST for 5min each wash, then rinsed with water, and 3’ phosphates on the ligated oligos were removed with 0.5U/ul Quick CIP in 1x CutSmart buffer by incubation at 4C 1hr under slow rotation followed by 37C 1hr. Samples were then again washed 3x in 2xSSCT, 5min per wash. Circ6G1 and Circ7G1 were prepared using T4 DNA ligase and short splint oligos^[a]^ (splint6F5 and splint7F5, respectively) and purified using a Zymo Oligo Concentrator spin column. Products were checked for size and purity via TBE-urea gel. CircDNA annealing mix (100nM Circ6G1, 100nM Circ7G1, in 1x hybridization buffer, containing 2x SSC, 10% formamide, 0.1% Tween-20) was added to samples and incubated overnight at 40C under slow rotation. Samples were then washed in hybridization buffer at 40C for 30min under slow rotation, and then washed in 2xSSCT, 1xSSCT, and finally 1xPBST, at 5min per wash.

#### Circular DNA annealing and RCA (rolling circle amplification)

Samples were rinsed with water and RCA mix (25ng/ul T4 gene 32, 1x phi29 reaction buffer, 0.5ug/ul rBSA, 250uM dNTP, 0.2U/ul phi29 polymerase) was added, incubated at 4C 1hr under slow rotation, and then 30C overnight. In cases where fluorescence was to be observed, RCA mix was supplemented with 20uM fluorescein-12-dUTP. Samples were washed 3x in 2xSSCT.

#### UEI oligos annealing and T4 DNA extension/ligation

UEI annealing mix (100nM 21.004G1/2-BC oligo mix, 100nM 21.073pt, 100nM 21.074B, 2xSSC, 5% formamide, 0.1% Tween-20) was added to samples, incubated at 4C 1hr under slow rotation, and then 50C 2hrs. After bringing to room temperature, samples were washed in UEI-hybridization buffer (2xSSC, 5% formamide, 0.1% Tween-20) 1hr at 50C, followed by washes in 2xSSCT, 1xSSCT, and then 1xPBST. After water rinse, extension/ligation mix (1x T4 ligase buffer including ATP, 1mM dNTP, 0.15U/ul T4 DNA polymerase, 20U/ul T4 DNA ligase) was added and incubated 1hr under slow ration at 4C, followed by room temperature incubation 40min. Samples were then washed 3x in PBST, followed by a water rinse.

#### IVT (In vitro transcription)

IVT-ligation mix was prepared by adding to final concentrations together, in order, oligo 21.075 (100nM), 21.066C3 (1uM), 1x IVT reaction buffer, 7.5mM rNTP mix, 100ng/ul T4g32, 0.5U/ul T4 RNA ligase 2, 0.25U/ul RppH, 10% T7 Enzyme Mix, 73.6ug/ul 4arm-PEG20K-Vinylsulfone, and 6.4ug/ul 3-arm Thiocure-333 (PEG reagents being thawed from -80C immediately prior to reaction). Mixes were added to individual zebrafish embryos at a total volume of 30ul. Hydrogel was allowed to form around samples for 2hrs at room temp. Reaction was then incubated at 37C 20hrs. Afterward, hydrogels were denatured via addition of 12ul denaturation solution (457.5mM KOH, 100mM EDTA, 42.5mM DTT) for 2hrs at 4C. Denaturation was stopped by addition of 12ul stop solution (600 mM Tris-HCl pH7.5, 0.4N HCl). After mixing, 30ul proteinase K mix (0.28% Tween-20, 0.09U/ul proteinase K, 8.6 mM Tris-HCl pH7.5) was added to the 54ul samples for a total of 84ul. This was incubated at 50C 1hr.

#### RNA isolation and cDNA synthesis

RNA was purified by addition of 1.2x RNAClean XP beads, following manufacturer protocols, and eluted into water. DNase I digestion was performed (final concentration of 0.8U/ul Superase-In, 0.1U/ul DNase I, 1x DNase I reaction buffer) at 37C for 30min. RNA was again purified via 1.2x RNAclean XP, and eluted into water. Reverse transcription was carried out in a final concentration of 500nM each of RT primers (21.077 and 21.085), 500uM dNTP, 1x FS buffer, 5mM DTT, 1U/ul Superase-In, and 10U/ul Superscript III. Primers and dNTP were added first to RNA/water-eluent and incubated at 65C 5min, after which the mixture was placed promptly on ice. The rest of the reaction mixture was then added, and samples were then incubated 1hr at 50C, followed by inactivation 15min at 70C, and kept at 4C. ExoI enzyme was then added directly to the product to final concentration of 3.3U/ul. After mixing, this was incubated at 37C for 30min, followed by heat-inactivation at 80C for 20min.

#### Library preparation

cDNA products (from IVT products) were then amplified in two separate PCR reactions. “cDNA-amplicons” were amplified by adding ExoI-digested product at a final 1:80 dilution into 4 separate reactions (per embryo) containing final concentrations of 300nM 21.046G1-BC primer, 300nM 21.081b primer, 1x HiFi PCR buffer, 200uM dNTP, 2mM MgSO_4_, and 0.02U/ul Platinum Taq HiFi. This reaction was thermocycled 95C 2min, 5x(95C 30s, 56C 30s, 68C 2min), 20x(95C 30s, 68C 2min), 68C 5min, 4C. “UEI-amplicons” were amplified by adding ExoI-digested product at a final 1:40 dilution into 2 separate reactions (per embryo) containing final concentrations of 300nM 21.077-G1 primer, 300nM 21.076BB primer, 3.3uM each of 4E4.interf1 and 4E.interf2 (3’P-capped oligos to interfere with PCR recombination^7;[b]^), 5% DMSO, 1x HiFi PCR buffer, 200uM dNTP, 2mM MgSO_4_, and 0.02U/ul Platinum Taq HiFi. This was thermocycled 95C 2min, 1x(95C 30s, 66C 30s, 68C 2min), 18x(95C 30s, 68C 2min), 68C 5min, 4C. PCR products were then purified using a 0.75x volume of Ampure XP beads, following manufacturer protocol. Products were quantified and sequenced on an Illumina NextSeq 500 instrument using 150-cycle kits (112nt read 1, 44nt read 2), including the sequencing primer sbs3b as a custom spike-in according to manufacturer protocol.

### Sequence Analysis

Sequence analysis was performed using the pipeline previously described^7^, with code updated for the larger scale of data available at https://github.com/wlab-bio/vdnamic.

Briefly, sequencing reads were demultiplexed via the barcodes depicted in Figure S1. Subsequently, for each amplicon type, sequence elements (UMI type I, UMI type II, UEIs, and cDNA inserts) were separately clustered using a 1bp difference-criterion using the EASL algorithm^7^.

For UEI data sets, each UEI was assigned a UMI-pair by plurality (relevant only if a specific UEI appeared to show two different pairings of UMIs – a signature of PCR recombination). The resulting “consensus pairings” were then pruned, with each UMI required to be associated with 2 UEIs, and each association (unique UMI-UMI pair) required to be associated by at least 2 reads. The largest contiguous matrix (found via single-linkage clustering, with rarefaction depicted in Figure 1J) was retained for image inference.

For cDNA insert sequence data, reads grouped by the same UMI had a sequence-consensus generated by majority-vote. These sequences were then trimmed to eliminate sequence adapters. Those inserts retaining at least 25bp of non-artificial sequences (at least among known artificial sequences) were then counted toward the cDNA-insert UMIs (depicted as rarefaction in Figure 1I). These consensus inserts were then inputted into STAR alignment^[c]^ using the Danio Rerio genome assembly GRCz11. Gene-assignments were performed using GTF annotations, with mappings ties (in edit-distance) between genes receiving equal weight, and priority assignment to rRNA in case of an ambiguous match with the genome.

The UMIs from genome-mapped cDNA-insert libraries (Table S1) were then matched back to the UMIs in the UEI amplicon libraries of the corresponding specimen. The gene-calls were then applied to label UMIs in the UEI-inferred image.

### Simulations

All simulations were performed by taking the raw coordinates depicted in Figs 2A,E and calculating Gaussian “point-spread functions”. For UMI *i* and UMI *j* at ground truth positions 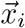 and 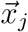, respectively, and with *N* being the sum-total of all counts in the simulated data set, we assigned a raw count *n*_*ij*_ *←* NegativeBinomial(mean = *μ*_*ij*_, *p*) where 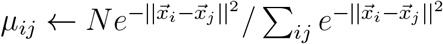 and *p* ← 0.8.

### Clustering

The preliminary segmentation analysis of GSE inferences (Figs 3H-I) was performed by taking UEI-associations collectively – and an equivalent total number of nearest neighbors (such that for *k* nearest neighbors, *k ← N*_UEIs_/*m*_UMIs_) – and calculating diffusion kernels within the GSE embedding coordinates 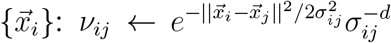 where 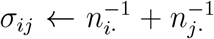. The symmetrized normalized Graph Laplacian matrix formed by the “pseudo-linkages” then underwent spectral clustering as previously described^7^ down to a conductance threshold of 0.2, requiring a minimum segment size of 50 UMIs.

Inter-segment UEI counts then defined a new UEI-count matrix that was row-normalized. The top eigenvector – specifically, its median – provided the boundary for memberships visualized in Figs 4A-B of cephalic vs caudal.

For gene-gene UEI matrices (Fig 4I), a similar summation of categories was performed as with segments above. The symmetrized normalized Graph Laplacian was used to generate 100 eigenvectors from which proximities to each of the molecular species, rRNA, MT-rRNA, and gDNA. For a gene *i* relative to any one of these molecular species, here designated *c*, this proximity was estimated through the Gaussian kernel 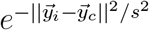 with 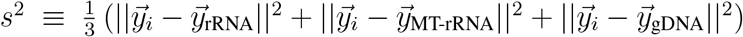. A linear transform was then applied to affix the locations of each rRNA, MT-rRNA, and gDNA to the vertices of each ternary plot.

**Table S1:** UMI/UEI statistics for embryos 1 and 2.

|  | Embryo 1 | Embryo 2 |
| --- | --- | --- |
| Total cDNA UMIs (with inserts >=25bp) | 8379427 | 8819189 |
| Total genome mapped cDNA-inserts | 5395011 | 5531597 |
| rRNA | 155559 | 222599 |
| MT-rRNA | 759518 | 682481 |
| Protein-coding/mRNA | 2946938 | 2885390 |
| Genomic (non-CDS) | 1436384 | 1654004 |
| Other | 96612 | 87123 |
| UEI-data matched cDNA-inserts | 1135105 | 1120439 |
| Total in UEI-matrix (largest contig) | 293352 | 300369 |
| Type I UMIs in UEI-matrix (largest contig) | 2118976 | 2252619 |
| Type II UMIs in UEI-matrix (largest contig) | 1644364 | 1776217 |
| Total UEIs in UEI-matrix (largest contig) | 8730431 | 9012629 |

**Table S2:**
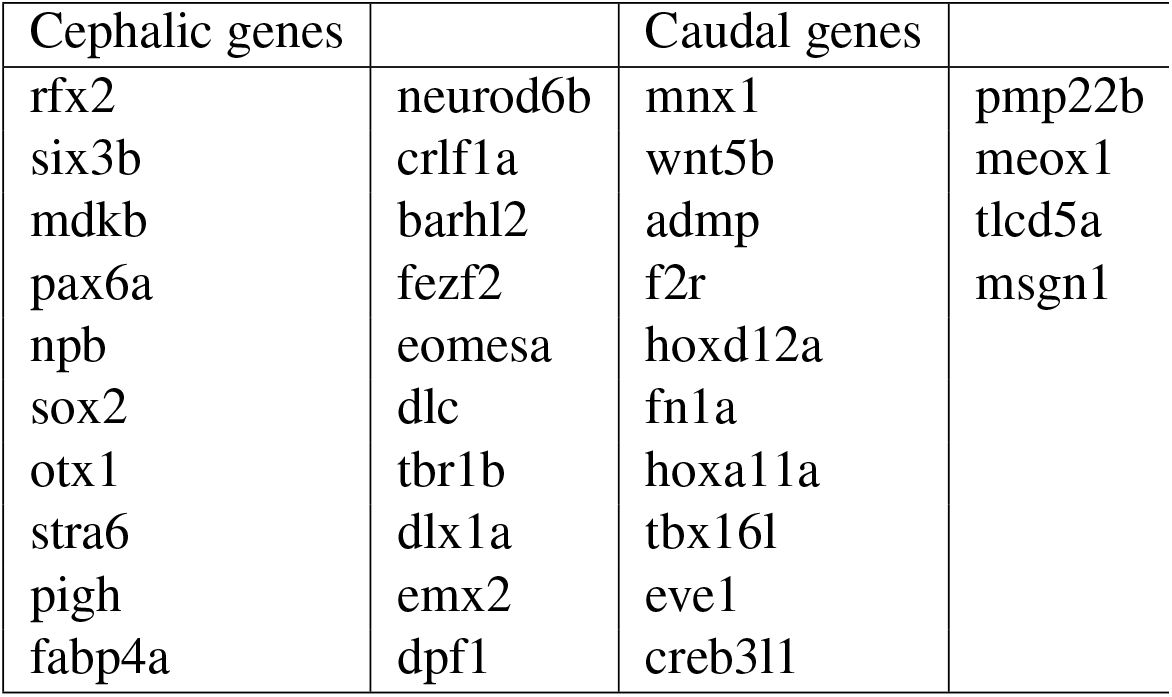
Cephalic and caudal genes used in Fig 4F-G. Cephalic genes were generated by performing a database search ^16;[d]^. Cephalic genes were collected by filtering for “telencephelon” and selecting those genes with clear evidence of predominant expression in the head in 24hpf embryos in ISH images. Caudal genes were collected by filtering for “caudal fin” and “tail bud” and selecting those genes with clear evidence of predominant expression in the caudal region in 24hpf embryos in ISH images.

### sMLE/UMAP/GSE comparisons

In Figs 2E-J, hyperparameters were chosen as follows. The top eigenvectors of the UEI matrix were applied to both sMLE (top 50, to undergo projected gradient descent) and UMAP (following common practice and to better constrain, the top 30 eigenvectors were used, with nearest neighbors set to 100). GSE used the top 50 eigenvectors (E = 50), along with 10 data tessellations. The total GSE eigenvectors used in the final projected gradients descent totaled 200.

### GSE (Geodesic Spectral Embeddings)

GSE begins by “de-indentifying” type I and type II UMIs in our data set (Fig S3A): taking the rectangular *m*_**I**_ *× m*_**II**_ UEI-count matrix of *m*_**I**_ type I UMIs and *m*_**II**_ type II UMIs and converting it into a square *m×m* symmetric matrix, consisting of *m* = *m*_**I**_ + *m*_**II**_ UMIs. UMI-UMI interactions are modeled statistically as illustrated in Fig S1B. There, the observed UEI matrix counts *n*_*ij*_ – associating UMI *i* with UMI *j* – are generated stochastically according to probabilities, *w*_*ij*_, that go up the closer UMI *i* is to UMI *j* in the embedding space and go down the further apart they are in the embedding space. The set of all UMI/point positions which collectively best comports with this model is what we will call the optimal GSE embedding.

Evaluating embedding positions in this way, however, most of all requires a way to estimate the distance between them, given the original count data. Most commonly, this is done by taking the subspace formed by the top *E* eigenvectors of the data matrix and finding a straight-line Euclidean distance between two points^[e][f][g]^ with the goal of identifying and focusing on nearest-neighbor relations. GSE does this as an initial processing step, but uses the resulting *putative* nearest neighbors only to form local tangent spaces about each point. These tangent spaces will allow us to estimate the *geodesic* distances along the d-dimensional surface (sitting in the full m-dimensional data space) we wish to represent in the final embedding.

These tangent spaces may involve the *E* “global” eigenvectors of the full symmetric count matrix in Fig S3A. However, global eigenvectors that describe the dominant axes of variance for the full data set may – on their own – be insufficient to differentiate the tangent spaces of neighboring points. To avoid this problem, GSE augments the global eigenvector subspace by performing several random, distinct tessellations of this subspace (Fig S3C). Each tessellation partitions the original points into 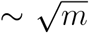 sectors of 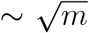 points each.

The choice of 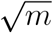 here is motivated by the fact that the total computational complexity of analyzing all sectors together will ultimately scale as the product of the number of tessellations and the number of points per tessellation, ie the total number of points in the data set 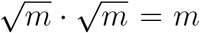. Each of these sectors now possesses smaller “local” count matrices generated by collapsing and summing the matrix elements belonging to all other sectors, as illustrated in Fig S3D. Each of these smaller matrices, because they include 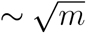 points and 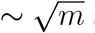 sectors, will be of size 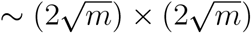.

GSE then appends the *E* eigenvectors generated from these local matrices to the original *E*-dimensional global eigenvector subspace, forming a fuller subspace spanned by 2*E* eigenvectors describing each point’s local neighborhood. These local neighborhoods are then used to find putative 2*E* nearest neighbors (the minimum to span the full eigenvector subspace) for each point in the data set (Fig S3E) using a standard kNN algorithm. The 2*E* nearest-neighbors for each point are then shuffled between tessellations, in order to allow each point – within each tessellation – to have a local neighborhood that extends beyond the boundaries of the sector into which it was assigned.

Although each sector now has a locally defined eigenvector subspace, this eigenvector subspace contains coordinates for the *collapsed* counts of all other sectors. These collapsed-sector coordinates can then be used to bridge the coordinates of points that have been assigned to different sectors (Fig S3F).

GSE then uses each point’s 2*E* nearest neighbors from before to perform a local PCA analysis, giving each point its own d basis vectors – where *d* is the low-dimensionality of the intended embedding – that constitute the local tangent spaces for the high-dimensional surface (Fig S4A). For point *i*, we use these basis vectors to construct a tangent space projection matrix **M**_*i*_. Projecting the original counts associating different data points onto these tangent spaces then allows calculation of *d×d* count covariance matrices **Σ**_*i*_ (Fig S4B). For any vector 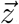, we can then write the re-scaled square-distance 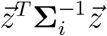 with a diffusion-distance metric 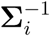 in the neighborhood of point *i*.

For any *given pair* of points *i* and *j* at positions 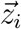 and 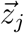, respectively, GSE uses the procedure described so far to estimate a geodesic distance between them in two steps. First, the shortest connecting path between 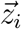 and 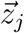 is estimated by adding difference-vectors projected onto their respective tangent spaces to give the intermediate points 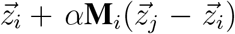 and 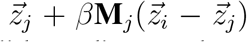. The scalar values *α* and *β* are then adjusted to minimize the Euclidean distance between the resulting vector sums (Fig S4C), which uniquely specifies a piecewise linear path from 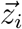 to 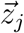. Second, the geodesic distance is estimated as the diffusion-distance traversed by this path, calculated using distance metrics 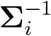 and 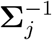 (Fig S4D).

GSE then collates these distances across all random tessellations (from Fig S3) and incorporates them together into a single geodesic similarity matrix that determines – for every point – the half of data points that are closer to it than the other half along the d-dimensional surface swept out by the tangent spaces calculated earlier (Fig S4E). GSE approximates this geodesic similarity matrix in a sparse matrix 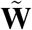 by, for every point *i*, randomly and uniformly selecting a set of other points across the data set, estimating their geodesic distances to point *i*, and retaining the lowest 1/2^*d*^ fraction of these distances. These retained points, along with the nearest-neighbors found in the original eigenvector subspace, are inserted into the corresponding row in the form of a sparse set of Gaussian proximities.

GSE uses the geodesic similarity matrix 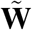 as part of what we call the “GSE matrix” 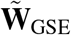: a mathematical description of how the original count matrix ought to be embedded across the *d*-dimensional tangent spaces used to construct 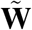 . The top eigenvectors of this matrix are considered to be putative solutions to the *d*-dimensional embedding problem. In serving this purpose 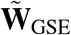 will be a projection of the count data *into* the geodesic similarity matrix so that 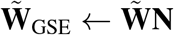.

Because we consider the top eigenvectors of 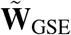 to fit the data to the global curvature of the data set, we now apply an incremental projected gradient descent on a global objective function (Fig S4F). The objective function we use here is the Kullback-Leibler divergence

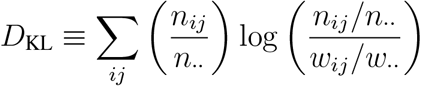

This is a statistical distance between the data counts *n*_*ij*_ and the proximities in the embedding space 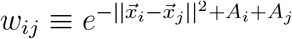, where 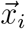 is the *d*-dimensional embedding coordinate of point *i* and where the amplitude 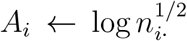. Here and elsewhere in this paper, subscript “·” denotes index summation, such that *n*_*i·*_ ≡ Σ_*j*_ *n*_*ij*_ .

Note that minimizing *D*_KL_ is equivalent to maximizing the log-likelihood ℒ of the multinomial that uses UEI-counts as “independent trials” on the space of all possible UMI-pairings – the framing in earlier DNA microscopy work^7^. This is because ℒ = Σ_*ij*_ *n*_*ij*_ *log w*_*ij*_/*w*_*‥*_ + constant, which is the same as minus-the-expression for D_KL_ once constant coefficients are factored out.

**Figure S1:**
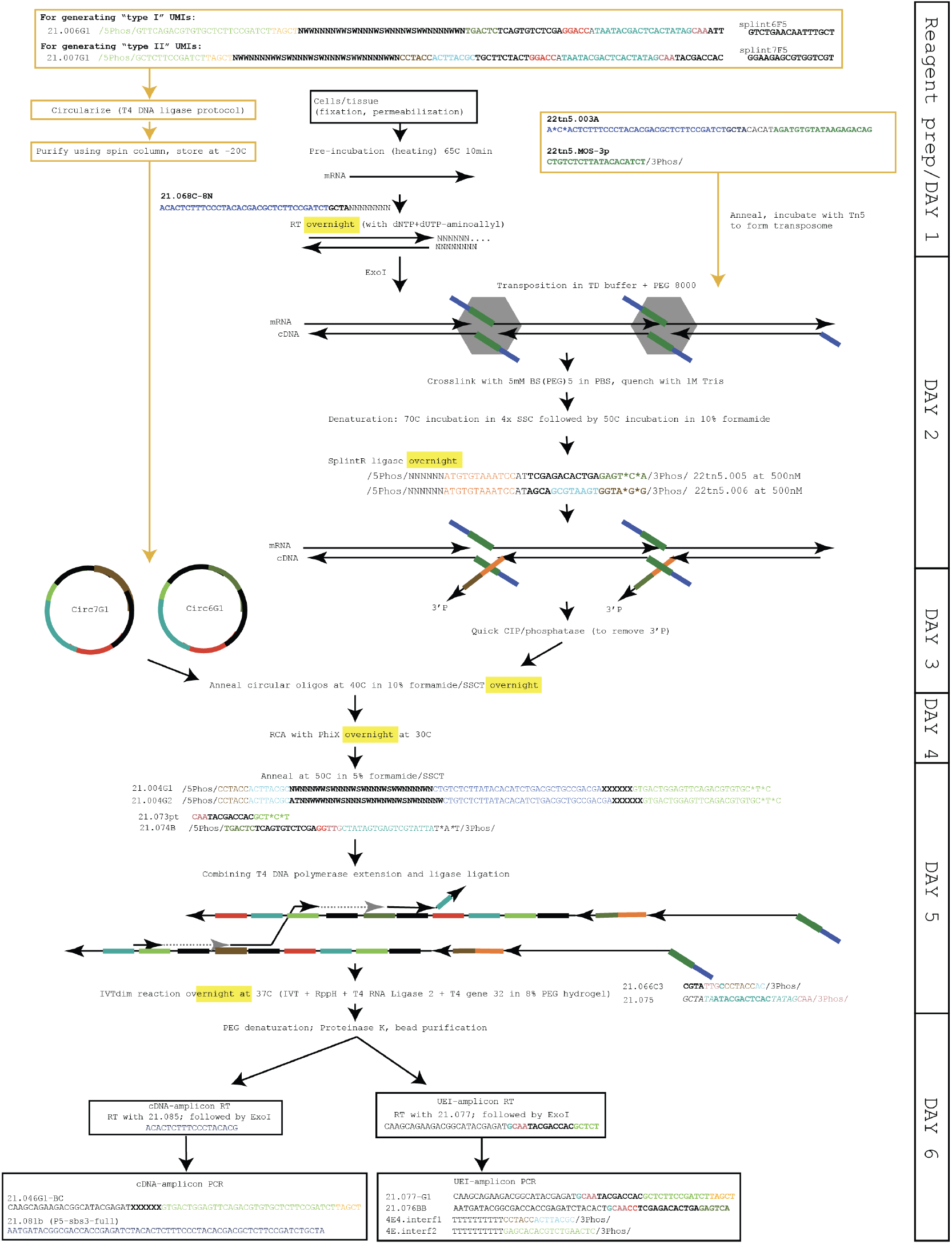
Experimental flow of volumetric DNA microscopy. “XXXXXX” corresponds to sample barcodes (arbitrary 6nt identifiers).

**Figure S2:**
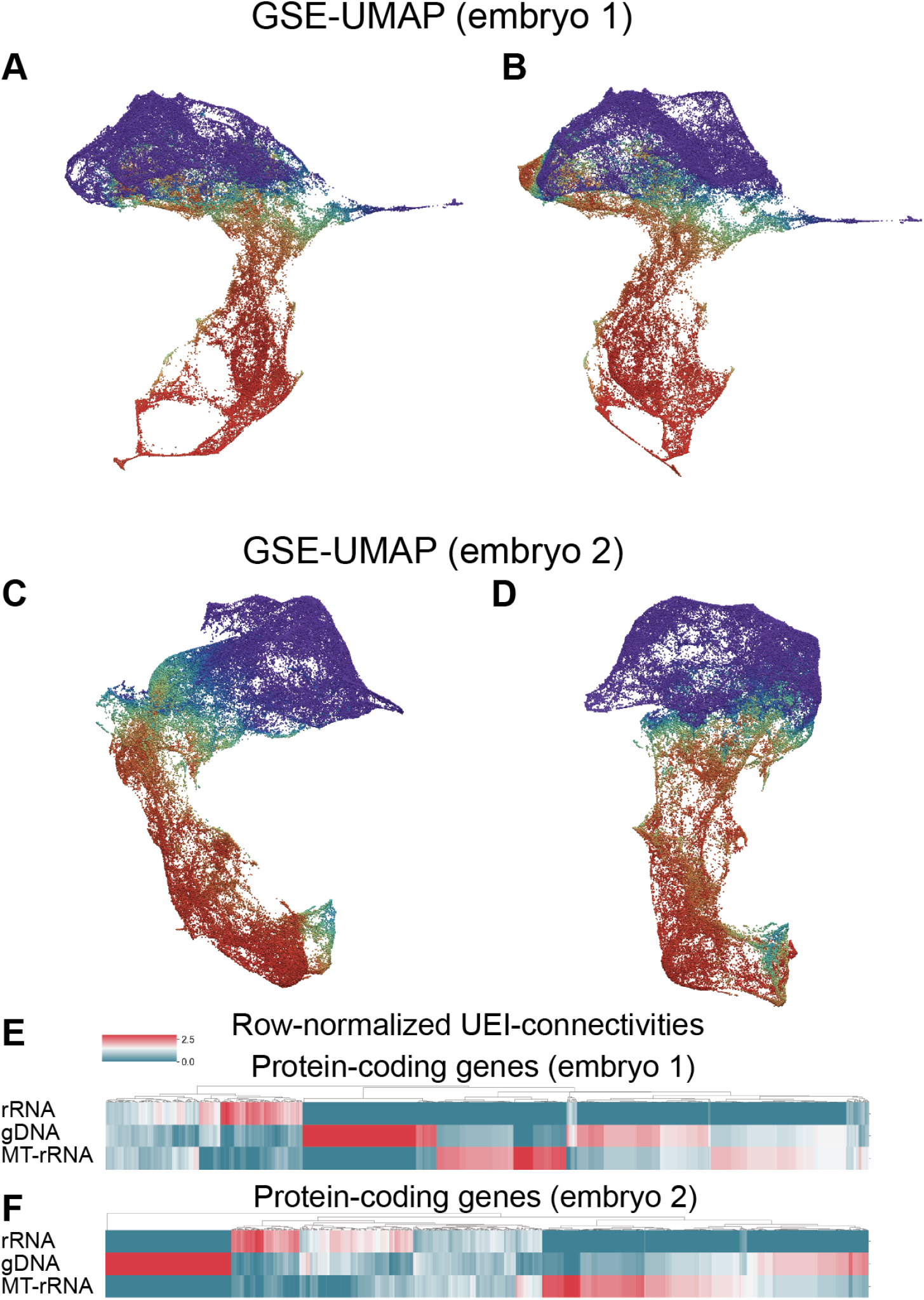
Spatio-genetic visualizations. GSE-UMAP plots from different perspective angles for embryos 1 (**A-B**) and 2 (**C-D**), with coloration as in Fig 4A-B,E. UEI connectivities in embryos 1 (**E**) and 2 (**F**) to molecular species rRNA, gDNA, and MT-rRNA, as in Fig 4H, but where UEI-counts are first normalized to the sum of each row, and are only then normalized to the mean for each protein-coding gene (columns).

**Figure S3:**
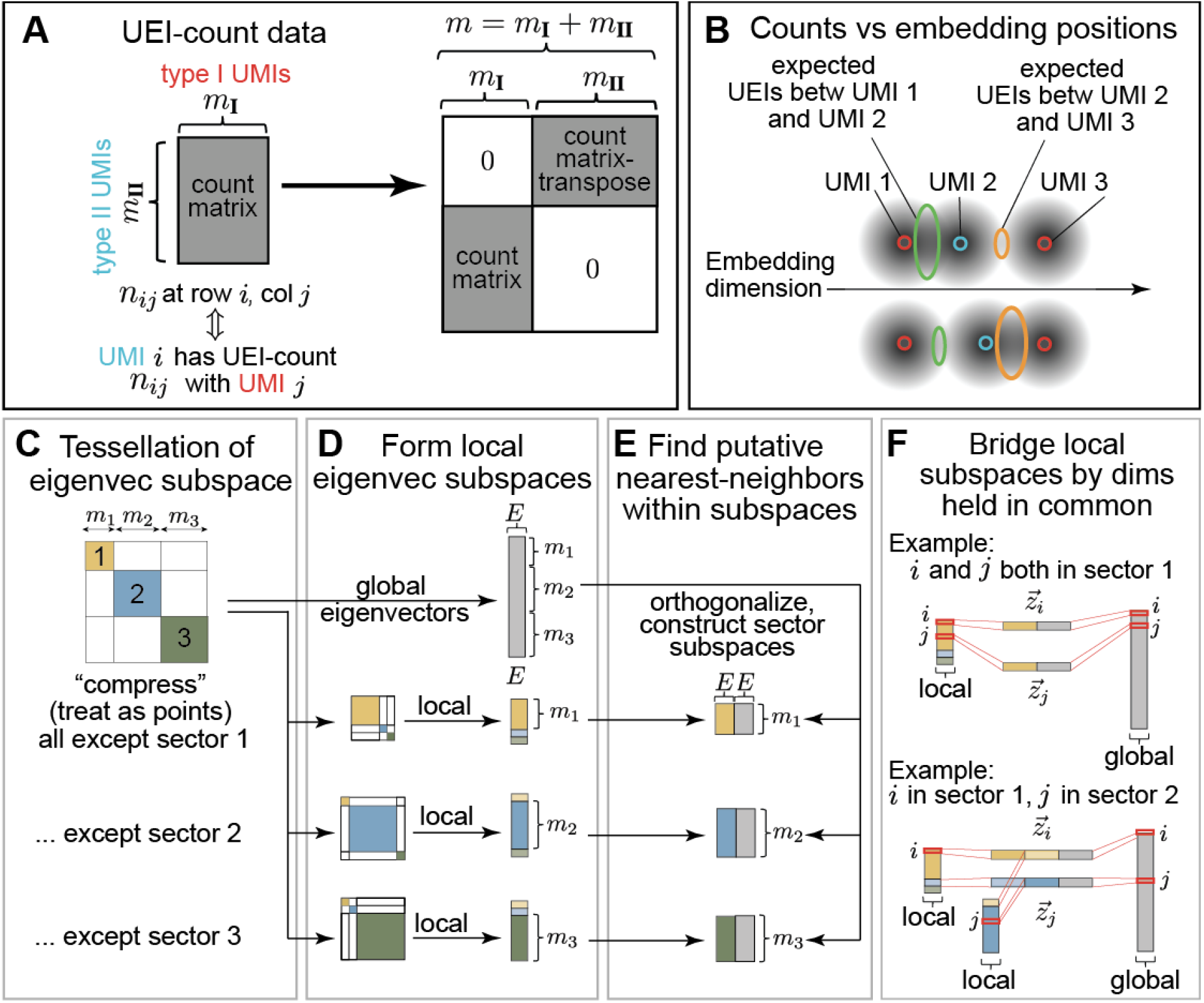
Assigning global and local subspace coordinates to UEI-count matrix observables. GSE begins by taking an arbitrary count matrix and transforming it into a block-symmetric matrix describing a bipartite graph (**A**). It asserts that the counts in this matrix describe the overlaps between diffusion “fields” of the rows and columns of this matrix in an embedding space (**B**). The GSE procedure begins by forming several random tessellations of the top eigenvectors of the *m×m* count matrix (**C**), with each tessellation consisting of multiple sectors (illustrated as sizes *m*_1_ + *m*_2_ + *m*_3_ = *m*). The top eigenvectors of each sector are calculated by collapsing the other sectors into single rows/columns of “local” count matrices (**D**). These eigenvector subspaces, combined with eigenvectors from the “global” count matrix, are used to calculate nearest neighbors on a per-sector/per-tessellation basis (**E**). The coordinates of any pair of points across the data set can then be compared by analyzing the eigenvector subspace dimensions they share (**F**).

**Figure S4:**
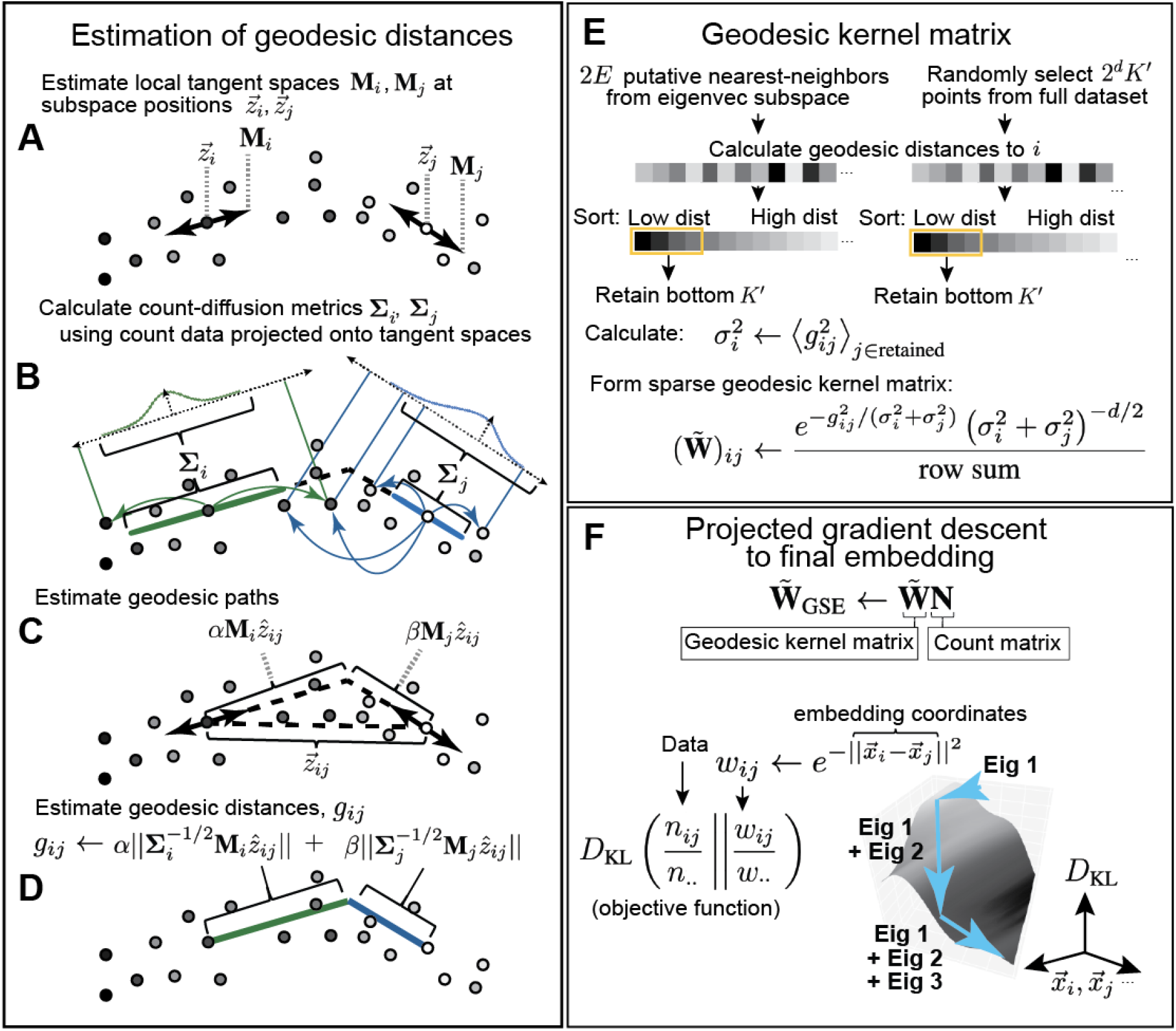
GSE’s numerical procedure to estimate the geodesic distance between any two points. begins by constructing local tangent spaces within the eigenvector subspaces in Fig S3 to construct local tangent spaces (**A**). Projecting each point’s counts onto its own local tangent space then allows the calculation of count covariance matrices, labeled as “count-diffusion metrics” (**B**). Then, taking any two points in the data set, a shortest piecewise-linear path is constructed using knowledge of tangent spaces alone (**C**). This path is then inputted into a rescaled distance that applies the count-diffusion metrics from earlier (**D**). This geodesic estimate can then be applied both to a point’s already established 2*E* nearest neighbors (from Fig S3) and to a random selection of 2^*d*^*K*^*′*^ other points – where here we set *K*^*′*^*←*10*d* – across the data set. Both sets of distances are sorted independently, and the lowest *K*^*′*^ distances from each set are retained and placed in a Gaussian kernel matrix (**E**). This kernel matrix is then used to generate a “geodesic kernel matrix”, 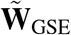, whose eigenvectors are then used to construct the solution to the embedding problem (**F**).

**TABLE S3.**

| Reagents | Manufacturer | Catalog number |
| --- | --- | --- |
| Pronase | Sigma-Aldrich | 10165921001 |
| Water | Invitrogen | 10977023 |
| PBS (10x, pH7.4) | Invitrogen | AM9624 |
| Paraformaldehyde (16%) | Thermo Scientific | 28906 |
| Methanol | Sigma-Aldrich | 34860-100ML-R |
| Tween-20 | Sigma-Aldrich | P9416-100ML |
| Thermolabile Proteinase K (0.12U/ul) | NEB | P8111S |
| Formamide | Sigma-Aldrich | 47671-250ML-F |
| rBSA (20 ug/ul) | NEB | B9200S |
| Superase-In (20 U/μL) | Invitrogen | AM2696 |
| dNTP | Thermo Scientific | R0181 |
| Aminoallyl-dUTP (50 mM) | Thermo Scientific | R1101 |
| SuperScript III Reverse Transcriptase (200 U/μL) | Invitrogen | 18080085 |
| ExoI (20U/ul) | NEB | M0293L |
| Tris-HCl (1M, pH8.0) | Fisher Scientific | BP1758-500 |
| NaCl (5M) | Invitrogen | AM9760G |
| Tagmentase (2ug/ul) | Diagenode | C01070010-20 |
| Glycerol | Sigma-Aldrich | G5516-100ML |
| Tagmentase dilution buffer | Diagenode | C01070011 |
| Tris-HCl (1M, pH7.5) | Fisher Scientific | BP1757-500 |
| MgCl<sub>2</sub> (1M) | Invitrogen | AM9530G |
| N,N-Dimethylformamide | Sigma-Aldrich | D4551-250ML |
| PEG8000 (50%) | NEB | B0216L |
| ATP (10mM) | NEB | B0216L |
| BS(PEG)5 | Thermo Scientific | A35396 |
| Tris (1M, pH8.0) | Invitrogen | AM9855G |
| SSC (20x) | Invitrogen | AM9770 |
| SplintR ligase (25U/ul) | NEB | M0375S |
| Quick CIP (5U/ul) | NEB | M0525S |
| Zymo Oligo Concentrator | Zymo Research | D4060 |
| TBE-urea Gels (15%) | Invitrogen | EC68855BOX |
| T4 gene 32 (10ug/ul) | NEB | M0300S |
| Phi29 polymerase (10U/ul) | NEB | M0269L |
| Fluorescein-12-dUTP (1mM) | Thermo Scientific | R0101 |
| T4 DNA polymerase (3U/ul) | NEB | M0203L |
| T4 DNA ligase (400U/ul) | NEB | M0202L |
| 4arm-PEG20K-Vinylsulfone | Sigma-Aldrich | JKA7025-1G |
| 3-arm Thiocure-333 | Bruno Bock | 345352-19-4 |
| T4 RNA ligase 2 (10U/ul) | NEB | M0239L |
| RppH (5U/ul) | NEB | M0356S |
| rNTP (25mM each) | NEB | N0466L |
| MEGAscript T7 Transcription Kit (T7 enzyme mix) | Invitrogen | AM1334 |
| KOH (1M) | Honeywell | 319376-500ML |
| EDTA (0.5M) | Sigma-Aldrich | 03690-100ML |
| DTT (1M) | Thermo Scientific | P2325 |
| HCl (1N) | Sigma-Aldrich | H9892-100ML |
| Proteinase K (0.8U/ul) | NEB | P8107S |
| RNAclean XP beads | Beckman Coulter | A63987 |
| DNase I (2U/ul) | NEB | M0303S |
| Platinum Taq HiFi (5U/μL) | Invitrogen | 11304029 |
| Ampure XP beads | Beckman Coulter | A63881 |
| Nextseq 500/550 Mid-Output v2.5 Kit (150 cycles) | Illumina | 20024904 |
| Nextseq 500/550 High Output Kit v2.5 (150 Cycles) | Illumina | 20024907 |

**TABLE S4.**
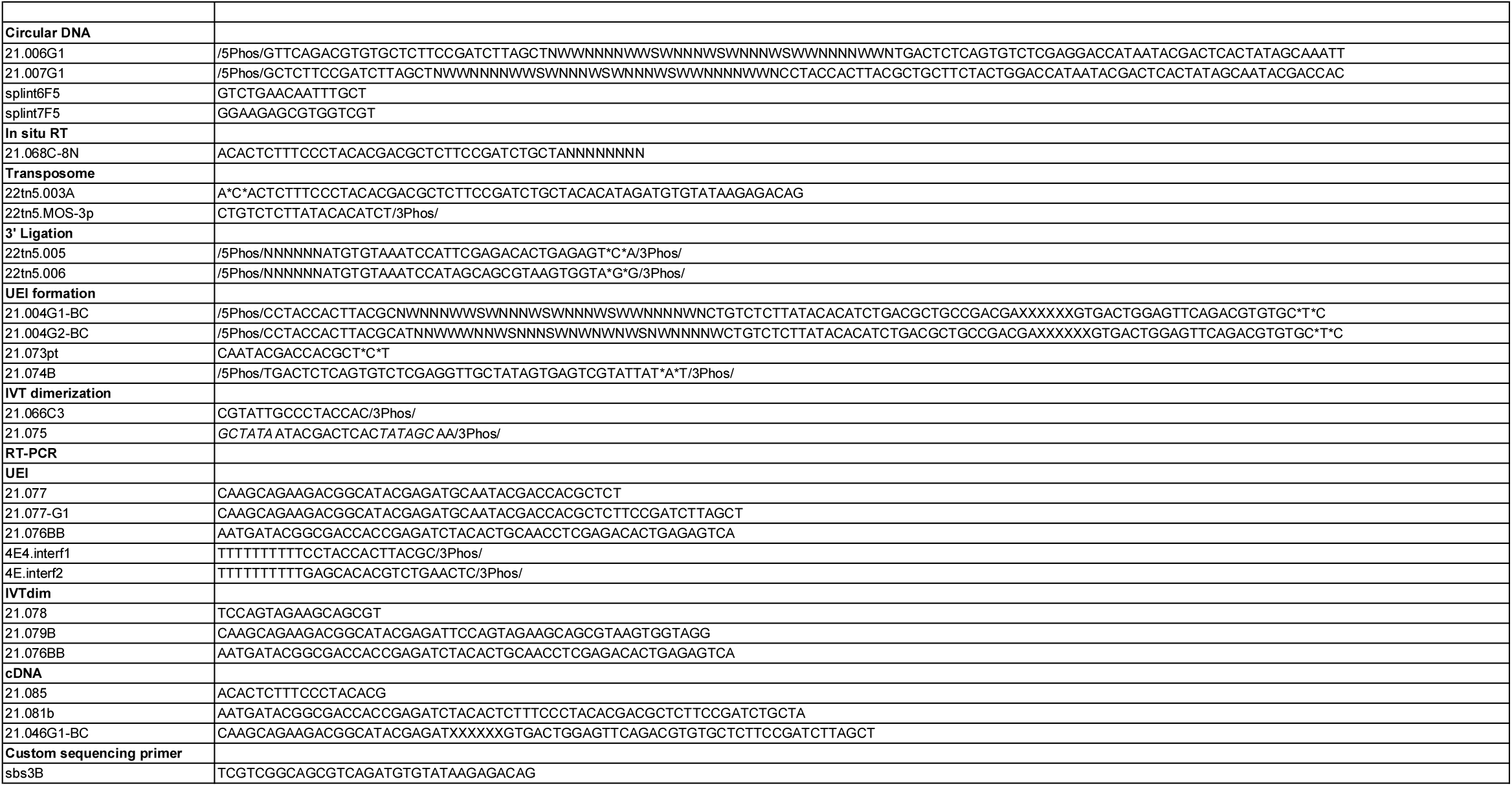

## Footnotes

[a] An, Ran, et al. Nucleic Acids Research 45.15 (2017): e139-e139.

[b] Turchaninova, Maria A., et al. European journal of immunology 43.9 (2013): 2507-2515.

[c] https://github.com/alexdobin/STAR

[d] ”The Zebrafish Information Network (ZFIN).” The Zebrafish Information Network, https://zfin.org.

[e] Coifman, Ronald R., and Stéphane Lafon. App and comp harmonic analysis 21.1 (2006): 5-30.

[f] Van der Maaten, Laurens, and Geoffrey Hinton. Journal of machine learning research 9.11 (2008)

[g] McInnes, Leland, John Healy, and James Melville. arXiv preprint arXiv:1802.03426 (2018).

## References

[1] Jeremy Thorpe, Ikeoluwa A Osei-Owusu, Bracha Erlanger Avigdor, Rossella Tupler, and Jonathan Pevsner. Mosaicism in human health and disease. Annual review of genetics, 54:487–510, 2020.

[2] Chenqu Suo, Krzysztof Polanski, Emma Dann, Rik GH Lindeboom, Roser Vilarrasa-Blasi, Roser Vento-Tormo, Muzlifah Haniffa, Kerstin B Meyer, Lisa M Dratva, Zewen Kelvin Tuong, et al. Dandelion uses the single-cell adaptive immune receptor repertoire to explore lymphocyte developmental origins. Nature Biotechnology, pages 1–12, 2023.

[3] Sara Bizzotto and Christopher A Walsh. Genetic mosaicism in the human brain: from lineage tracing to neuropsychiatric disorders. Nature Reviews Neuroscience, 23(5):275–286, 2022.

[4] Lisa N Waylen, Hieu T Nim, Luciano G Martelotto, and Mirana Ramialison. From whole-mount to single-cell spatial assessment of gene expression in 3d. Communications biology, 3(1):602, 2020.

[5] Anjali Rao, Dalia Barkley, Gustavo S França, and Itai Yanai. Exploring tissue architecture using spatial transcriptomics. Nature, 596(7871):211–220, 2021.

[6] Shahar Alon, Daniel R Goodwin, Anubhav Sinha, Asmamaw T Wassie, Fei Chen, Evan R Daugharthy, Yosuke Bando, Atsushi Kajita, Andrew G Xue, Karl Marrett, et al. Expansion sequencing: Spatially precise in situ transcriptomics in intact biological systems. Science, 371(6528):eaax2656, 2021.

[7] Joshua A Weinstein, Aviv Regev, and Feng Zhang. Dna microscopy: optics-free spatio-genetic imaging by a stand-alone chemical reaction. Cell, 178(1):229–241, 2019.

[8] Filip Karlsson, Tomasz Kallas, Divya Thiagarajan, Max Karlsson, Maud Schweitzer, Jose Fernandez Navarro, Louise Leijonancker, Sylvain Geny, Erik Pettersson, Jan Rhomberg-Kauert, et al. Molecular pixelation: Single cell spatial proteomics by sequencing. bioRxiv, June 2023.

[9] Ian T Hoffecker, Yunshi Yang, Giulio Bernardinelli, Pekka Orponen, and Björn Högberg. A computational framework for dna sequencing microscopy. Proceedings of the National Academy of Sciences, 116(39):19282–19287, 2019.

[10] Alexander A Boulgakov, Erhu Xiong, Sanchita Bhadra, Andrew D Ellington, and Edward M Marcotte. From space to sequence and back again: Iterative dna proximity ligation and its applications to dna-based imaging. bioRxiv, page 470211, 2018.

[11] Liyi Xu, Ilana L Brito, Eric J Alm, and Paul C Blainey. Virtual microfluidics for digital quantification and single-cell sequencing. Nature methods, 13(9):759–762, 2016.

[12] Xiao Wang, William E Allen, Matthew A Wright, Emily L Sylwestrak, Nikolay Samusik, Sam Vesuna, Kathryn Evans, Cindy Liu, Charu Ramakrishnan, Jia Liu, et al. Three-dimensional intact-tissue sequencing of single-cell transcriptional states. Science, 361(6400):eaat5691, 2018.

[13] Etienne Becht, Leland McInnes, John Healy, Charles-Antoine Dutertre, Immanuel WH Kwok, Lai Guan Ng, Florent Ginhoux, and Evan W Newell. Dimen-sionality reduction for visualizing single-cell data using umap. Nature biotechnology, 37(1):38–44, 2019.

[14] Olivier Armant, Martin März, Rebecca Schmidt, Marco Ferg, Nicolas Diotel, Raymond Ertzer, Jan Christian Bryne, Lixin Yang, Isabelle Baader, Markus Reischl, et al. Genome-wide, whole mount in situ analysis of transcriptional regulators in zebrafish embryos. Developmental biology, 380(2):351–362, 2013.

[15] Mark van Eekelen, John Overvoorde, Carina van Rooijen, and Jeroen den Hertog. Identification and expression of the family of classical protein-tyrosine phosphatases in zebrafish. PLoS One, 5(9):e12573, 2010.

[16] Bernard Thisse,S Pflumio, M Fürthauer, B Loppin, V Heyer, A Degrave, R Woehl, A Lux, T Steffan, XQ Charbonnier, et al. Expression of the zebrafish genome during embryogenesis. ZFIN direct data submission, 2001.

